# A defect in COPI-mediated transport of STING causes immune dysregulation in COPA syndrome

**DOI:** 10.1101/2020.05.20.106500

**Authors:** Zimu Deng, Zhenlu Chong, Christopher S. Law, Kojiro Mukai, Frances O. Ho, Tereza Martinu, Bradley J. Backes, Walter L. Eckalbar, Tomohiko Taguchi, Anthony K. Shum

**Affiliations:** Department of Medicine, University of California San Francisco, San Francisco, CA; Laboratory of Organelle Pathophysiology, Department of Integrative Life Sciences, Graduate School of Life Sciences, Tohoku University, Sendai, Japan; Toronto Lung Transplant Program, University Health Network, University of Toronto, Toronto, ON, Canada; Cardiovascular Research Institute, University of California San Francisco, San Francisco, CA

## Abstract

Pathogenic *COPA* variants cause a Mendelian syndrome of immune dysregulation with elevated type I interferon signaling^1,2^. COPA is a subunit of coat protein complex I (COPI) that mediates Golgi to ER transport^3^. Missense mutations that disrupt the COPA WD40 domain impair binding and sorting of proteins targeted for retrieval to the ER but how this causes disease remains unknown^1,4^. Given the importance of COPA in Golgi-ER transport, we speculated that type I interferon signaling in COPA syndrome involves missorting of STING. Here we show that a defect in COPI transport due to mutant COPA causes ligand-independent activation of STING. Furthermore, SURF4 is an adapter molecule that facilitates COPA-mediated retrieval of STING at the Golgi. Activated STING stimulates type I interferon driven inflammation in *Copa^E241K/+^* mice that is rescued in STING-deficient animals. Our results demonstrate that COPA maintains immune homeostasis by regulating STING transport at the Golgi. In addtion, activated STING contributes to immune dysregulation in COPA syndrome and may be a new molecular target in treating the disease.

---

COPA syndrome is a genetic disorder of immune dysregulation caused by missense mutations that disrupt the WD40 domain of COPA^1^. COPA is a subunit of coat protein complex I (COPI) that mediates retrograde movement of proteins from the Golgi apparatus to the endoplasmic reticulum (ER)^3^. Prior studies have shown that alterations to the COPA WD40 domain lead to impaired binding and sorting of proteins bearing a Carboxyl-terminal dilysine motif as well as a defect in retrograde Golgi to ER transport^1,4^. To date, the molecular mechanisms of COPA syndrome remain unknown including whether missorted proteins are critical for initiating the disease.

A clue to the pathogenesis of COPA syndrome recently arose with the observation that type I interferon signaling appears to be highly dysregulated in the disease^2^. This led us to investigate whether COPA syndrome shares features with any of the well-described Mendelian interferonopathy disorders^5^. COPA syndrome manifests similarly to the type I interferonopathy STING-associated vasculopathy with onset in infancy (SAVI). Both diseases present at early age with interstitial lung disease, activation of type I interferon stimulated genes (ISG) and evidence of capillaritis^6,7^. STING is an ER-localized transmembrane protein involved in innate immune responses to cytosolic nucleic acids. After binding cyclic dinucleotides STING becomes activated as it translocates to the ER-Golgi intermediate compartment (ERGIC) and Golgi. At the ERGIC/Golgi, STING forms multimers and activates the kinase TBK1 which subsequently phosphorylates the transcription factor IRF3 to induce expression of type I interferons and other cytokines^8^. In SAVI, gain of function mutations cause STING to aberrantly exit the ER and traffic to the Golgi and become activated^9^. Prior work has suggested that COPI may be involved in STING transport at the Golgi, but this is not well established and the molecular interactions between COPI and STING remain unknown^8,10^. Because COPA plays a critical role in mediating Golgi to ER transport, we hypothesized that activation of type I interferon signaling in COPA syndrome involves missorting of STING.

To examine this, we assessed lung fibroblasts from a COPA syndrome patient to determine if there was evidence of STING activation. We measured mRNA transcript levels of several interferon stimulated genes and found they were significantly elevated in comparison to healthy control lung fibroblasts in the presence or absence of a STING agonist (Fig. 1a and Extended Data Fig. 1a, b). Confocal microscopy of COPA syndrome fibroblasts revealed prominent colocalization of STING with the ERGIC (Fig. 1b). Western blots of cellular protein lysates showed an increase in STING multimerization (Fig. 1c) consistent with localization of STING at the ERGIC/Golgi and also higher levels of phosphorylated TBK-1 (pTBK1) and phosphorylated STING (pSTING) (Fig. 1d, Extended Data Fig. 1c), indicative of STING activation^11^. These data suggest that the elevated type I ISGs observed in COPA syndrome patients may be caused by spontaneous activation of STING.

**Fig. 1.**
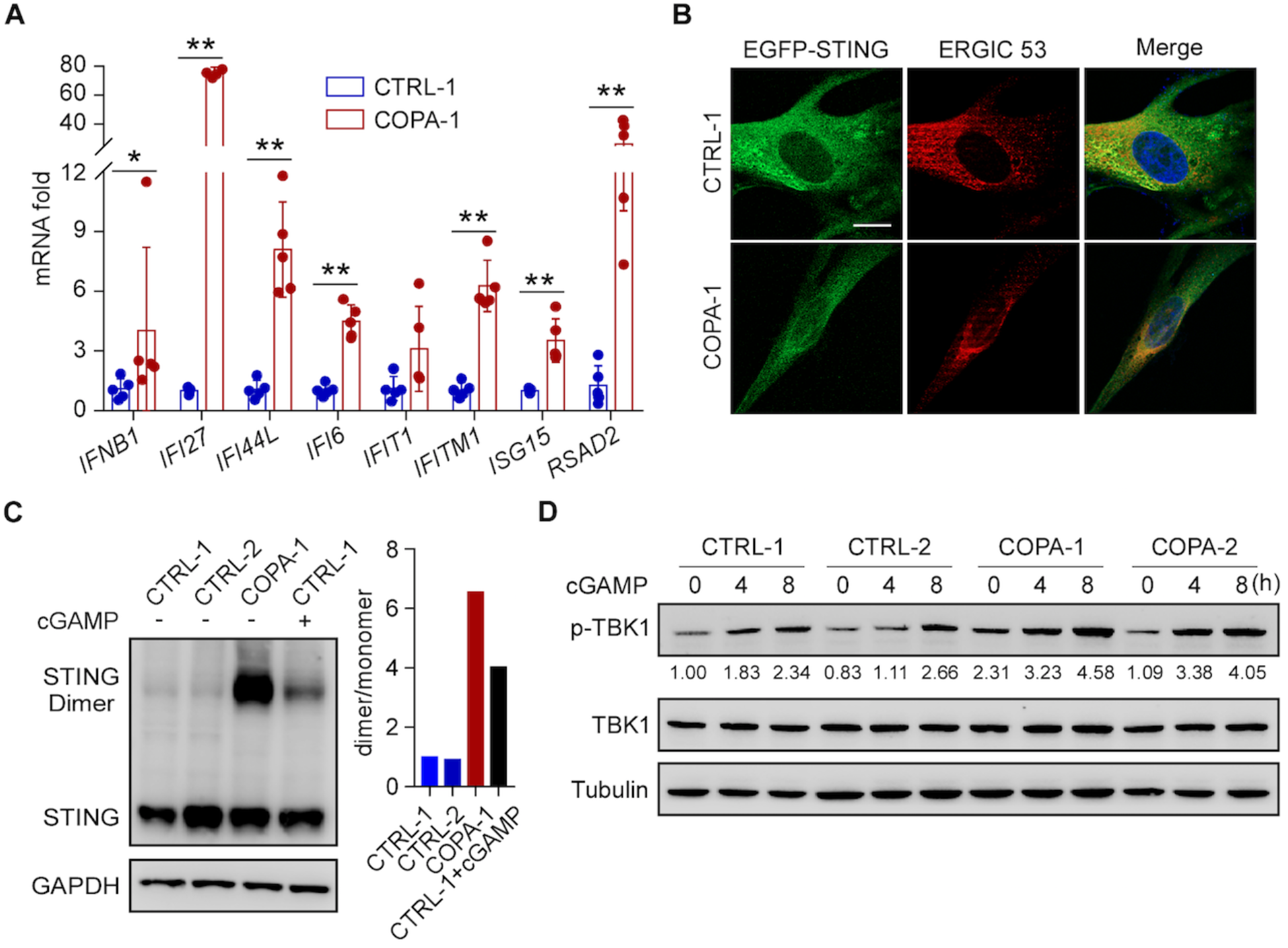
COPA syndrome patient fibroblasts demonstrate spontaneous STING activation. (**A**) Real-time PCR was performed for ISG expression in primary lung fibroblasts from a healthy control and COPA syndrome subject. (**B**) Primary lung fibroblasts from a healthy control and COPA syndrome subject were reconstituted with wild-type EFGP-STING retrovirus. The reconstituted cells were stained with indicated antibody and localization of STING observed using a confocal microscope. ERGIC 53 is an ERGIC marker. (**C**) Immunoblots of endogenous STING in primary lung fibroblasts from 2 healthy controls and a COPA syndrome subject under nonreducing SDS-PAGE conditions with and without cGAMP (0.5μg/ml) stimulation. (**D**) Immunoblots of indicated antibodies in primary lung fibroblasts with cGAMP (2μg/ml) stimulation for indicated time (*n*=2 per group). Data in (A) from two independent experiments represent means ± SD. *p<0.05, **p<0.01, (two-tailed Mann-Whitney *U* test). Scale bar, 20μm in (B).

To establish the specific role of mutant COPA in triggering STING pathway activation, we transduced cells with retroviral vectors encoding EGFP-STING and then transfected them with plasmids encoding wild-type or E241K mutant COPA. We performed confocal microscopy to examine whether mutant COPA caused STING to localize on the ERGIC/Golgi, similar to what we observed in patient fibroblasts. In cells expressing wild-type COPA, EGFP-STING was normally distributed throughout the cytoplasm^8^ whereas in cells with E241K mutant COPA, EGFP-STING co-localized with the Golgi marker GM130 (Fig. 2a). We next reconstituted HEK293T cells (which lack endogenous STING) with retroviral vectors encoding EGFP-STING and then transfected cells with wild-type or E241K mutant COPA. We found that even in the absence of a STING agonist, cells demonstrated a significant increase in pTBK1 (Fig. 2b) and higher levels of mRNA transcripts encoding *IFNB1* and type I ISGs (Fig. 2c). Importantly, all of these increases were abolished in HEK293T cells without EGFP-STING, indicating that mutant COPA activates TBK1 and type I interferon signaling specifically through STING rather than other innate immune pathways such as TRIF or MAVS (Fig 2b, c)^12^. Taken together, these data show that mutant COPA causes ligand-independent activation of STING on the Golgi with upregulation of type I interferon signaling.

**Fig. 2.**
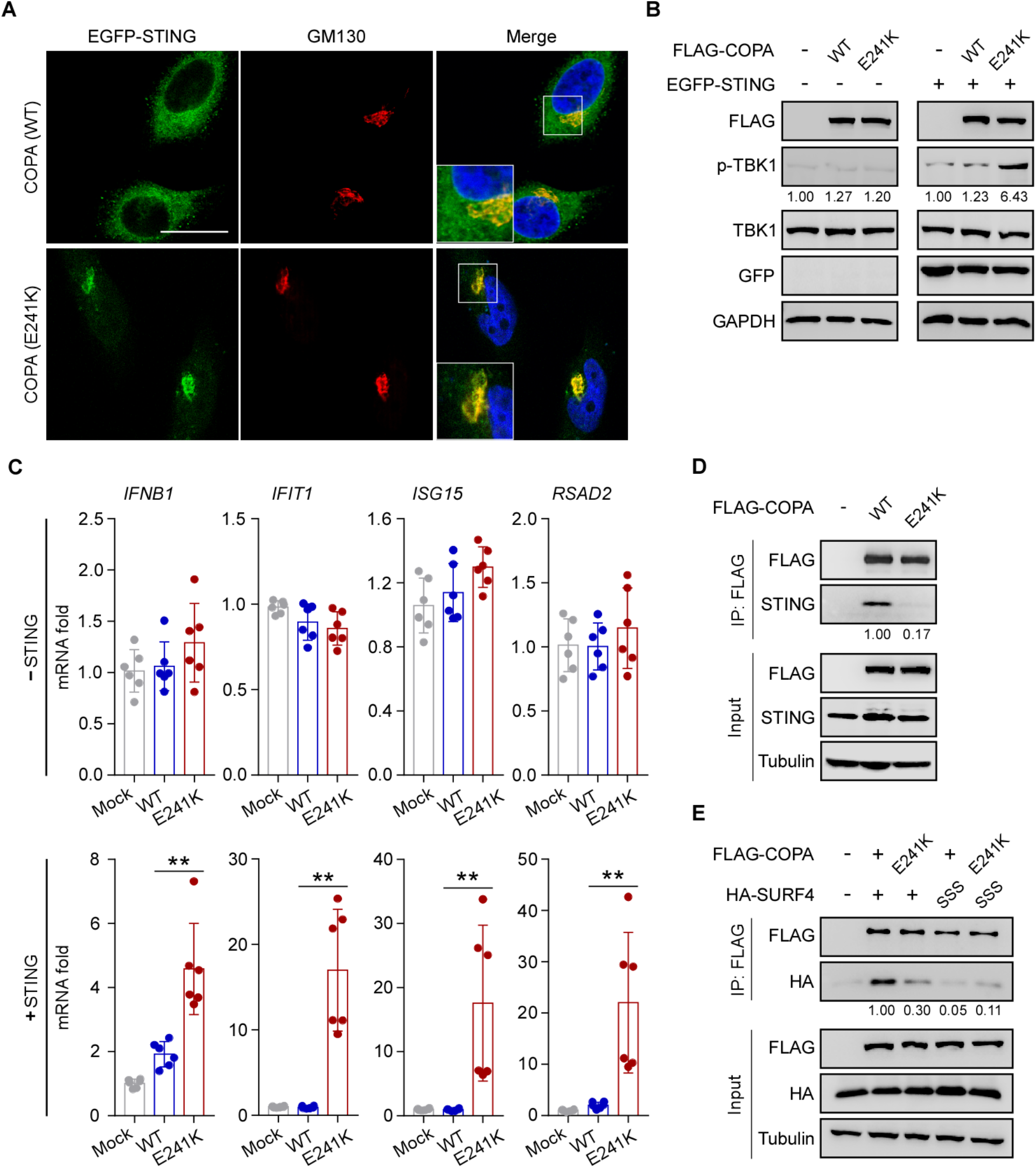
Activated STING is retained on the Golgi after failed retrieval of SURF4 by mutant COPA. (**A**) HeLa cells reconstituted with wild-type EGFP-STING retrovirus and transfected with wild-type COPA or mutant COPA (E241K) for 24 hours. Cells were stained with indicated antibodies and localization of STING observed using a confocal microscope. GM130 is a Golgi marker. (**B**) HEK293T cells with and without EGFP-STING were transfected with wild-type or mutant COPA (E241K) for 24 hours and then immunoblot performed with indicated antibodies. (**C**) HEK293T cells with and without EGFP-STING were transfected with wild-type or mutant COPA (E241K) for 24 hours and then real-time PCR performed for ISG expression. (**D**) FLAG immunoprecipitates (IP) from lysates of HEK293 cells overexpressing FLAG-tagged COPA (WT/E241K) were immunoblotted for indicated antibodies. (**E**) FLAG immunoprecipitates from lysates of HEK293T cells overexpressing FLAG-tagged COPA (WT/E241K) and HA-tagged SURF4 (WT/mutant) immunoblotted for indicated antibodies. Data in (C) from 2 independent experiments represent means ± SD. **p<0.01 (two-tailed Mann-Whitney *U* test). Scale bar, 20μm in (A).

We hypothesized that retention of STING on the Golgi might reflect a failure of STING to be taken up into mutant COPA containing COPI complexes for transport to the ER. To evaluate this, we analyzed the protein-protein interaction between COPA and STING. We performed coimmunoprecipitation experiments and found that although we could pull-down STING with wild-type COPA the amount of STING that co-immunoprecipitated with E241K mutant COPA was substantially reduced (Fig. 2d). We previously showed that disease-causative *COPA* mutations cause a defect in binding between the COPA WD40 domain and C-terminal dilysine motif (e.g. KKxx and KxKxx) of proteins targeted for retrieval to the ER^1^. Because STING lacks a dilysinetag, we hypothesized that an adaptor protein mediates the interaction between STING and COPA. A review of published studies uncovered seven STING interacting partners that contain a C-terminal dilysine motif (Extended Table 1)^13–16^. Among these, SURF4 was the only protein shown to cycle between the ER and Golgi via COPI^3,17^ and function as a cargo receptor^18^. Thus, we speculated that SURF4 was a likely candidate for mediating an interaction between COPA and STING. We confirmed through co-immunoprecipitation assays that SURF4 associates with STING and COPA (Fig. 2e and Extended Data Fig. 2a) and then mutated the C-terminal dilysine motif of SURF4 by replacing the lysines at positions −3, −4 and −5 from the C-terminus with serines (SURF4-SSS). The amount of SURF4-SSS pulled down with COPA was significantly less in comparison to wild-type SURF4, demonstrating the importance of the dilysine motif to the SURF4-COPA interaction (Fig. 2e). Consistent with this, we found that the association between SURF4 and mutant COPA was markedly reduced in comparison to wild-type COPA, reflecting a defect in binding between the mutant COPA WD40 domain and SURF4 di-lysine tag (Fig. 2e). Finally, as further evidence that SURF4 functions as an adapter molecule for STING and COPA, loss of SURF4 led to a reduction in the amount of STING that co-immunoprecipitated with wildtype COPA (Extended Data Fig. 2b) and also led to an increase in transcript levels of *Ifnb1* and type I ISGs (Extended Data Fig. 2 c). In aggregate, our data suggests that SURF4 functions as a cargo receptor for STING and that mutant COPA is unable to bind SURF4 and incorporate STING into COPI vesicles. This results in retention of STING on the Golgi where it becomes spontaneously activated and triggers type I interferon signaling.

We next turned to a mouse model of COPA syndrome to understand how mutant COPA-mediated STING activation causes immune dysregulation *in vivo.* We previously reported that *Copa^E241K/+^* mice which express one of the same disease causative mutations as patients spontaneously develop activated cytokine-secreting T cells and T cell-mediated lung disease^19^. Through bone marrow chimera and thymic transplant experiments, we showed that mutant COPA within thymic epithelial cells perturbs thymocyte development and leads to a defect in immune tolerance. Because STING is highly expressed in thymic tissue^20^, we wondered how missorting of STING due to mutant COPA might contribute to the T cell phenotypes we observed in *Copa^E241K/+^* mice.

To evaluate this, we first confirmed that *Copa^E241K/+^* mice demonstrate retention of activated STING on the Golgi with elevated type I interferon signaling, similar to what we observed in patient cells and our transfection assays (Fig. 3a-c). We next performed bulk RNA sequencing to identify gene expression programs that were altered in thymic epithelial cells. Genes involved in response to viruses were highly enriched in the set of genes differentially expressed between *Copa^E241K/+^* and wild-type mice in medullary thymic epithelial cells (mTEC) (Extended Data Fig. 3). Among the differentially expressed genes, we found significant upregulation of *Ifnb* and several type I ISGs, consistent with STING activation (Fig. 4a). Thymocytes migrating through the thymic epithelium were impacted by higher *Ifnb* levels because they exhibited an elevated type I interferon signature (Fig. 4b) and higher levels of Qa2 (Fig. 4c), a cell surface marker expressed in response to IFN-β^21^. We examined thymocyte populations using a staining protocol that subsets increasingly mature single positive (SP) cells into semi-mature (SM), mature 1 (M1) and mature 2 (M2) populations (Fig. 4d). Prior studies found that type I interferons are required during the transition of thymocytes from SM to M2 cells^21^. In *Copa^E241K/+^* mice, we found a significant increase in the proportion of M2 cells (Fig. 4d), suggesting that higher IFN-β levels promoted an expansion of late stage thymocytes. To determine the role of STING in these changes, we crossed *Copa^E241K/+^* mice to STING-deficient *Sting1^gt/gt^* mice. We measured mRNA transcripts of thymic epithelial cells for *Ifnb1* and type I ISGs and found that all of the increases returned to wild-type levels in *Copa^E241K/+^xSting1^gt/gt^* mice (Fig. 4e). In addition, loss of STING abrogated all of the thymocyte phenotypes caused by mutant COPA including the elevated type I IFN signature, increased Qa2 expression and higher proportion of M2 cells (Fig. 4c-e). Taken together, these data indicate that activated STING in the thymus of *Copa^E241K/+^* mice perturbs T cell development by elevating type I interferons to promote late stage thymocyte maturation.

**Fig. 3.**
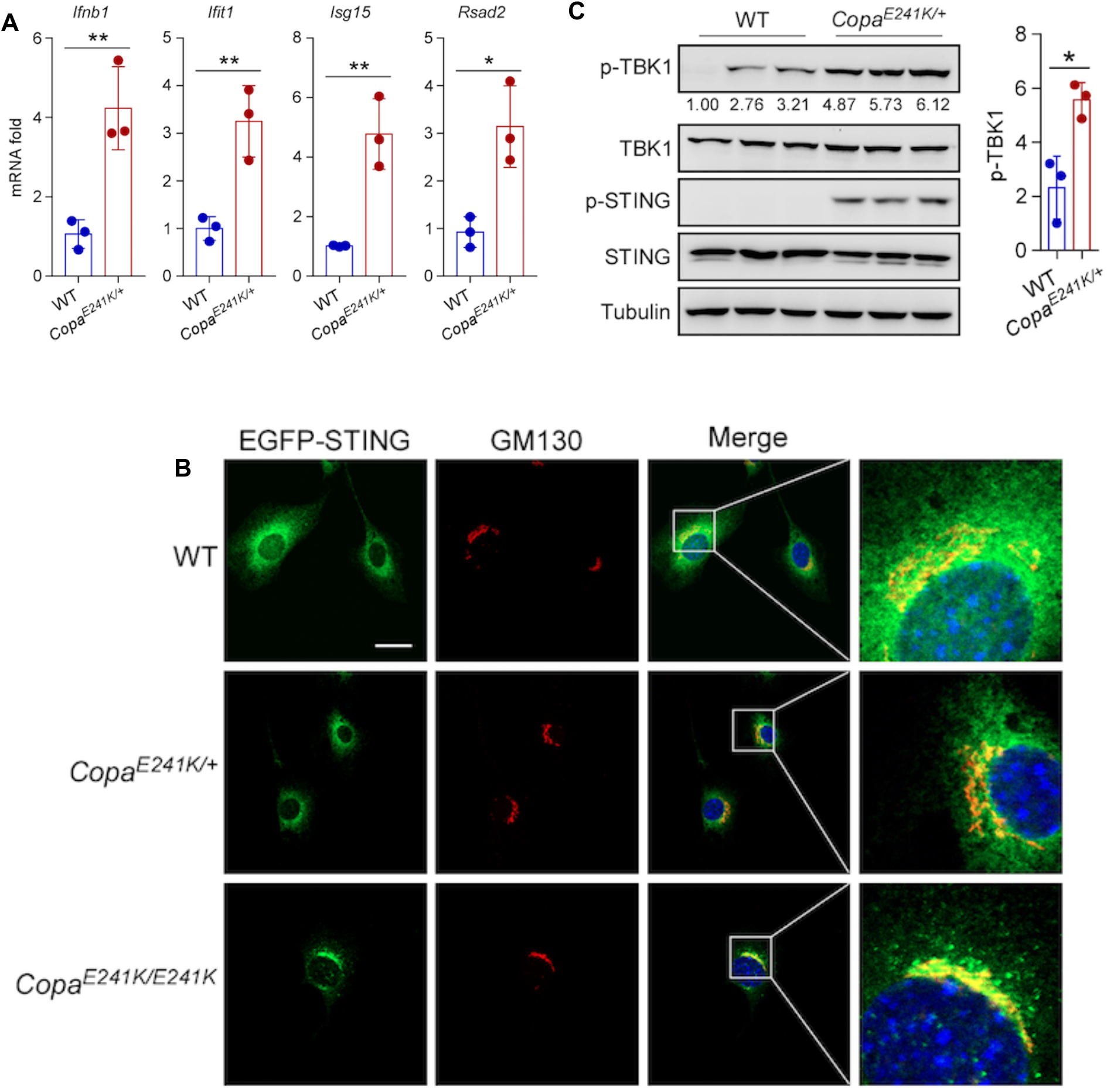
*Copa^E241K+^* mice exhibit STING activation with elevated type I interferon signaling. (**A**) Real-time PCR was performed for ISG expression in immortalized MEF cells derived from 3 WT and 3 *Copa^E241K/+^* mice. (**B**) Immortalized MEF cells from WT, *Copa^E241K/+^* and *Copa^E241K/IĪ24EK^* mice were reconstituted with wild-type EGFP-STING retrovirus. The reconstituted cells were stained with the indicated antibodies and localization of STING observed using a confocal microscope. GM130 is a Golgi marker. (**C**) Immunoblot was performed with the indicated antibodies in splenocytes from WT or *Copa^E241K/+^* mice (*n*=3 for each genotype). Data in (A, C) were means ± SD. *p<0.05, **p<0.01(unpaired two-tailed Student’s *t* tests). Scale bar, 20μm in (B).

**Fig. 4.**
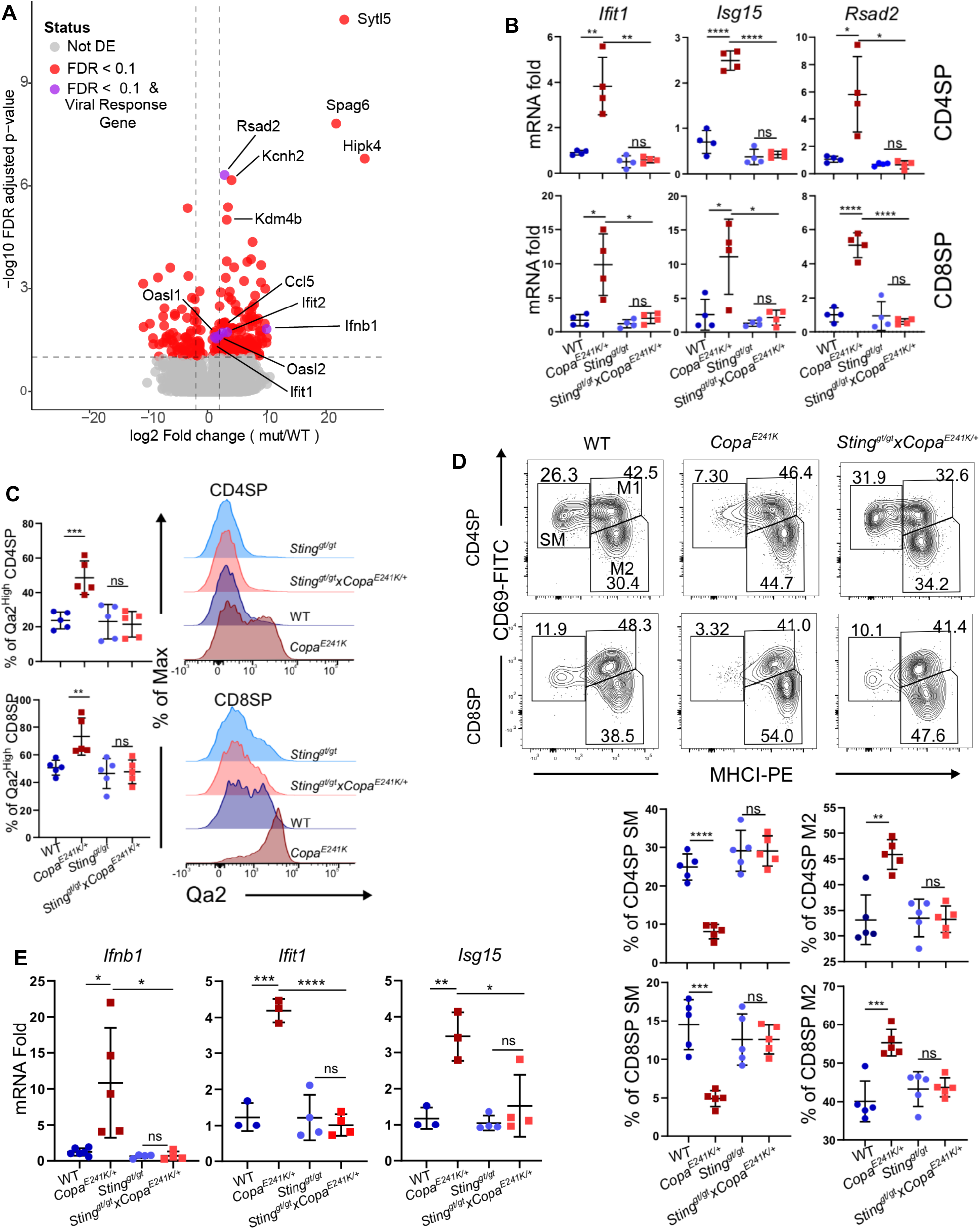
Activated STING perturbs thymocyte development by increasing type I interferons in the thymus. (**A**) Medullary thymic epithelial cells sorted from wild-type and *Copa^E241K^* mice were used for RNA-seq analysis (*n*=3 each genotype). Volcano plot of the differentially affected genes (FDR<0.1, red; FDR<0.1, ISG from viral response GO categories, purple). (**B**) Real-time PCR performed for *Ifit1, Rsad2* and *Isg15* expression in SP thymocytes (top:CD4SP; bottom:CD8SP). (*n*=4 mice each genotype, 2 independent experiments). (**C**) left: Percentages of Qa2^high^ population among SP thymocytes (top: CD4SP; bottom: CD8SP). right: Representative flow analysis of Qa2 expression on SP thymocytes (top: CD4SP; bottom: CD8SP). (*n*=5 each genotype, 3 independent experiments). (**D**) top: CD69 versus MHCI expression on SP thymocytes (top:CD4SP; bottom:CD8SP). bottom: Percentages of SM and M2 population among SP thymocytes (top: CD4SP; bottom: CD8SP). (WT: *n*=5 each genotype, 3 independent experiments). (**E**) Real-time PCR performed for *Ifnb1, Ifit1* and *Isg15* expression in thymic epithelial cells (CD45^-^, Epcam^+^,Ly51^-^,MHC-II^high^mTEC) (*n*≥3 each genotype, 2 independent experiments). Data in (B-F) represent means ± SD. *p<0.05, **p<0.01, ***p<0.001, ****p<0.0001, ns, not significant (unpaired, parametric, two-tailed student’s *t*-test).

We next focused on peripheral lymphoid organs to examine whether mutant COPA-mediated STING-activation contributed to systemic inflammation in *Copa^E241K/+^* mice. We reasoned that if STING were to have a significant role in systemic disease, this would provide an opportunity to test whether targeting STING activation dampens systemic inflammation in COPA syndrome patients. Splenocytes from *Copa^E241K/+^* mice had a significant increase in type I ISGs that completely reverted to wild-type levels in *Copa^E241K/+^xSting1^gt/gt^* mice (Extended Data Fig. 4a). An analysis of peripheral T cell populations revealed that STING deficiency reversed the significant increase in activated effector memory cells and cytokine-secreting T cells caused by mutant COPA (Fig. 5a and Extended Data Fig. 4b-d). Finally, in strong support of STING being a critical mediator of disease in COPA syndrome pathogenesis, we found that loss of STING rescued embryonic lethality of homozygous *Copa^E241K/E241K^* mice (Extended Table 2). In aggregate, these data indicate that STING contributes to immune dysregulation in *Copa^E241K/+^* mice and that targeting STING in COPA syndrome may be an important therapy for patients.

**Fig. 5.**
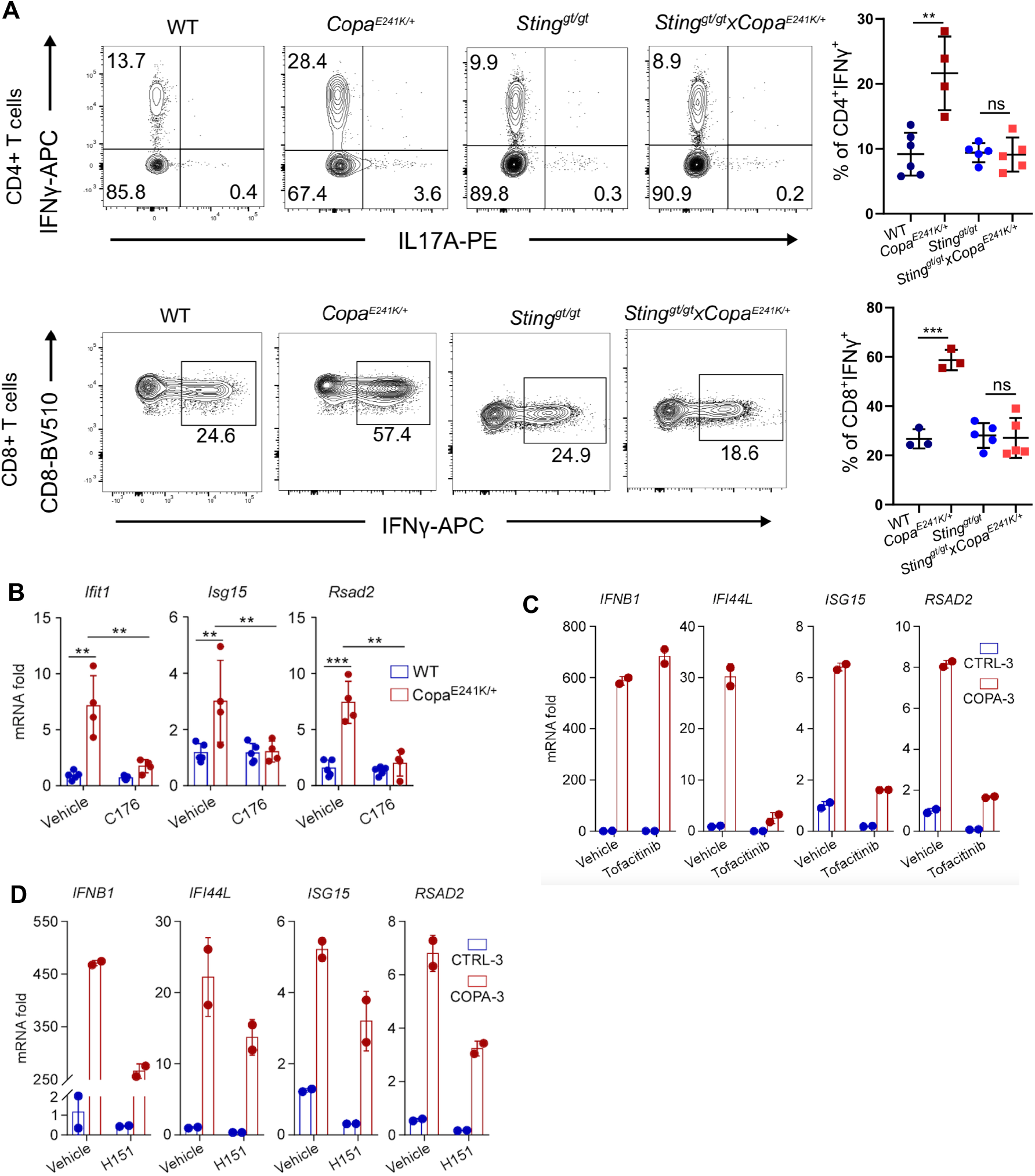
Loss of STING function dampens inflammation caused by mutant COPA. (**A**) Top left: intracellular levels of IFNγ and IL17A in splenic CD4+ T cells after PMA/Ionomycin stimulation. Top right: percentages of IFNγ producing CD4+ T cells. (*n*≥3 each genotype, >3 independent experiments). Bottom left: intracellular IFNγ production in splenic CD8+ T cells after PMA/Ionomycin stimulation. Bottom right: percentages of IFNγ producing CD8+ T cells. (*n*≥3 each genotype, >3 independent experiments). (**B**) Real-time PCR for ISG expression in splenocytes harvested from 4 WT mice and 4 *Copa^E241K/+^mice* treated with or without STING inhibitor C-176 (10μM) for 24 hours. (**C**) Real-time PCR performed in duplicate for ISG expression in PBMCs from a healthy control and COPA syndrome subject treated with or without STING inhibitor H-151 (10μM) or (**D**) JAK inhibitor Tofacitinib (2μM) for 24 hours. Data represent means ± SD. **p<0.01, ***p<0.001, ns, not significant. (unpaired, parametric, two-tailed Student’s t tests).

Activation of STING requires palmitoylation at the Golgi^22^ and recent studies report that inhibition of activation-induced palmitoylation of STING with small molecules prevents multimerization of the protein and recruitment of downstream signaling partners^23^. We treated splenocytes from *Copa^E241K/+^* mice with C-176 mouse STING inhibitor and found that this was highly effective at dampening activation of type I ISGs (Fig. 5b). We then took peripheral blood mononuclear cells (PBMCs) from a COPA syndrome patient or healthy control subject and treated them with the small molecule human STING inhibitor H-151. In parallel, we treated PBMCs with a JAK-STAT inhibitor since drugs in this class have recently been reported as a clinical therapy for COPA syndrome^24^. Although both the JAK-STAT inhibitor and STING inhibitor reduced type I ISGs, only the STING inhibitor was able to cause a decrease in the levels of the upstream cytokine IFN-β (Fig. 5c, d). Our results suggest that targeting STING activation with small molecule inhibitors might be an effective treatment for COPA syndrome patients with increased efficacy over current therapies.

This work establishes COPA as a critical regulator of STING transport that maintains immune homeostasis by mediating retrieval of STING from the Golgi. Defects in COPA function cause STING to multimerize and become spontaneously activated at the Golgi even in the absence of STING ligand, suggesting that at steady state low levels of STING continuously cycle between the ER and Golgi. In support of our findings, Mukai et. al used cultured cells to show that retrograde transport by COPI is essential to maintaining STING in its dormant state independent of cGAMP synthase (cGAS)^28^. Although we used a candidate approach to identify SURF4 as a putative cargo receptor that engages STING for incorporation into COPI vesicles by COPA, Mukai et. al directly tested 18 STING-binding proteins containing C-terminal dilysine motifs and found that knockdown of SURF4 was the only one that caused STING to localize on the Golgi^28^. Taken together, our data demonstrates an important role for SURF4 in mediating the retrieval of STING from the Golgi by COPI and provides new insight into the steady state dynamics of STING transport in resting cells.

Missense mutations in *COPA* that lie in the WD40 domain impair retrograde transport of a broad range of COPI cargo proteins containing C-terminal dilysine motifs^1^. Despite this, our data suggests that impaired trafficking of STING in particular is key to the pathogenesis of COPA syndrome, since not only does loss of STING reverse many of the immunological derangements in our mouse model, it also strikingly rescues embryonic lethality of homozygous mutant mice. Further study should be undertaken to determine whether other COPA cargo contribute to COPA syndrome pathogenesis independent of STING.

The ubiquitous expression of COPA has made it difficult to identify the cell types that are responsible for causing disease particularly since COPA is not enriched in any specific immune or lung cell. The tissue specificity of STING may provide additional insight to understanding COPA syndrome and the organs most affected. One unexpected outcome of our work was the finding that STING has a functional role in thymic stromal tissue. Although thymic epithelial cells are known to be a significant source of type I interferons^21,25^, the mechanisms regulating interferon secretion in the thymus remain largely unexplored. Interestingly, STING also modulates autophagy^8^ which in thymic epithelial cells is essential for processing self-antigen peptides for presentation to thymocytes^26^. Additional research might address whether activated STING alters autophagic function in thymic epithelial cells and if so, whether this impacts T cell selection.

Our work indicates that COPA syndrome belongs to a category of diseases defined by STING activation^5^. Going forward, clinicians may want to compare and contrast clinical features and treatment approaches in COPA syndrome and SAVI to establish optimal care of patients. Understanding the mechanisms by which STING causes interstitial lung disease has the potential to substantially improve outcomes in both disorders^6,7^. For those with COPA syndrome, the dramatic reversal of immune dysregulation we observed in *Copa^E241K/+^xSting1^gt/gt^* mice provides some hope that small molecule STING inhibition can be an effective molecularly targeted approach for treating this highly morbid disease.

## Methods

### Reagents

2’3’-cGAMP and CP-690550 (Tofacitinib) were purchased from InvivoGen. STING inhibitors N-(4-iodophenyl)-5-nitrofuran-2-carboxamide (C176) and 1-(4-ethylphenyl)-3-(1H-indol-3-yl)urea (H151) were synthesized by SYNthesis med chem at or greater than 99% purity by HPLC.

### Plasmids

Human STING and SURF4 were subcloned from HEK293T cDNA into pCMV6-AC (Origene) with a FLAG tag at the C-terminus or a HA tag at the N-terminus, respectively. Plasmids expressing FLAG tagged wild-type and mutant E241K human COPA were previously generated^1^. Retroviral plasmids expressing EGFP tagged human and mouse STING were kindly provided by Dr. Tomohiko Taguchi (Tohoku University).

### Study subjects

Subjects were selected on the basis of COPA syndrome diagnosis, with their written informed consent, were studied via protocols approved by the Research Ethics Board of Toronto General Hospital and the Institutional Review Boards for the protection of human subjects of Cleveland Clinic or the University of California San Francisco.

### Isolation of human lung fibroblasts

Control lung tissue was harvested from anonymous brain-dead donors from the Northern California Transplant Donor Network. Screening criteria for selection of healthy lung for specimen collection were previously described^2^. Mutant lung explants were from two COPA syndrome patients receiving lung transplants. Fibroblasts were isolated as described^3^. In brief, tissue was minced with scissors into 5 mm pieces and digested three times with 0.25% trypsin (GE Healthcare Life Sciences) for 10 minutes at 37°C. The resulting cell and tissue suspension was collected, neutralized with complete media (45% Ham’s F12, 45% DMEM, 10% FBS), pelleted, plated onto FBS coated 60 mm dishes and cultured in 5% CO_2_ at 37°C. Fibroblasts were ready to passage and use one week following isolation.

### Isolation of mouse embryonic fibroblasts

MEFs were isolated as described^4^. In brief, day 13.5 embryos were washed with PBS, minced with scissors into 1-2 mm pieces and digested three times with 0.25% trypsin for 10 minutes at 37°C. The cell suspension was neutralized with complete media (DMEM, 10% FBS), pelleted, resuspended in complete media, and cultured in 5% CO_2_ at 37°C.

To create immortal MEFs, Phoenix packaging cells (kindly gifted by Dr. Mark Anderson, UCSF) were transfected with pBABE-neo largeTcDNA plasmid (a gift from Bob Weinberg; Addgene plasmid # 1780; http://n2t.net/addgene:1780; RRID:Addgene_1780)^5^, and viral supernatant was collected two days later. Early passage MEFs were incubated in viral supernatant for two days and then transformed cells were selected with G418 (Teknova).

### siRNA knockdown

Predesigned siRNA oligomers (ON-TARGETplus SMARTPool) for *SURF4* and transfection control (siGENOME RISC-Free) were obtained from Dharmacon and resuspended in RNase-free water at 10 μM. siRNAs were transfected in Opti-Mem media (Life Technologies) for 48 h with Lipofectamine RNAiMAX (Life Technologies) according to manufactuer’s protocol.

### Immunoblotting and antibodies

Cells were lysed in Cold Spring Harbor NP40 lysis buffer (150 mM NaCl, 50 mM Tris pH 8.0, 1.0% Nonidet-P40) supplemented with protease and phosphatase inhibitors (PMSF, NaF, Na3VO4, Roche PhosSTOP) and then centrifuged at 12,000g for 15 mins at 4°C to get cellular lysate. Equal amounts of protein were loaded and size separated on an SDS-PAGE gel and wet transfered onto PVDF membrane. The membrane was blocked in TBS-T buffer with 5% milk for 1 h at room temperature, followed by overnight incubation with primary antibodies diluted in TBS-T with 5% BSA. Membrane was washed 3 times with TBS-T buffer for 10 mins, incubated at room temperature with HRP-conjugated IgG secondary antibody (Jackson Immunoresearch), washed 3 times with TBS-T for 10 mins followed once with TBS buffer for 10 mins, and then developed with SuperSignal West Femto Chemiluminescent Substrate (Life Technologies).

Rabbit antibodies against TBK1, p-TBK1 (Ser172), STING, p-STING (Ser365), p-STING (Ser366), FLAG, HA and GFP were from Cell Signaling Technology. Rabbit antibody against SURF4 was from Novus. Mouse antibodies against GAPDH and β-Tubulin were from Santa Cruz Biotechnology. Mouse antibody against GM130 was from BD Biosciences.

### Co-immunoprecipitation

HEK293 or HEK293T cells transfected with indicated plasmids were collected, washed once with PBS, lysed in NP40 lysis buffer supplemented with protease and phosphatase inhibitors and centrifuged at 12,000g for 15 min at 4°C. The supernatant was mixed with FLAG-M2 beads (Sigma) and incubated overnight at 4°C. Ten percent of lysate was saved as input. The following day, the beads were washed 3 times with IP washing buffer (10 mM Tris, pH 7.4, 1 mM EDTA, 150 mM NaCl and 1% Triton X-100) for 10 mins. 2× SDS loading buffer (100 mM Tris pH 6.8, 4% SDS, 20% glycerol, 10% β-mercaptoethanol, 0.1% bromophenol blue) was added into the IP complex and boiled at 95 °C for 5 min. Samples were analyzed by immunoblot as described above.

### Confocal microscopy

Cells were seeded onto glass coverslips and treated as indicated. Cells then were fixed with 4% paraformaldehyde, permeabilized with 0.1% Triton X-100 and blocked with 3% BSA. Slides were incubated with primary antibody overnight at 4°C, incubated with fluorescent-conjugated secondary antibody for 1 h at room temperature and followed with DAPI incubation for 10 mins at room temperature. Slides were then mounted with FluorSave Reagent (Millipore) and kept at 4°C in the dark. Images were captured with a Leica TCS SPE microscope.

Mouse antibody against ERGIC-53 was from Santa Cruz Biotechnology.

### Mice strains

*Copa^E241K^* knock in mice were generated in our lab^6^. C57BL/6J-*Sting1^gt^*/J (Sting^gt/gt^) mice were purchased from the Jackson Laboratory. All mice were maintained in the specific pathogen free (SPF) facility at UCSF, and all protocols were approved by UCSF’s Institutional Animal Care and Use Committee (IACUC).

### Flow cytometry and antibodies

Single cell suspension of thymocytes and splenocytes were prepared by mechanically disrupting the thymus and spleen. Cells were filtered through 40 μm filters (Genesee Scientific) into 15 ml conical tubes and maintained in RPMI 1640 containing 5% FBS on ice. Splenocytes were further subjected to red blood cells lysis (Biolegend). For evaluation of surface receptors, cells were blocked with 10 μg/ml anti-CD16/32 for 15 minutes at room temperature and then stained with indicated antibodies in FACS buffer (PBS, 2% FBS) on ice for 40 minutes.

For intracellular cytokine detection, freshly isolated splenocytes were stimulated by PMA (Sigma) and ionomycin (Sigma) in the presence of brefeldin A (Biolegend) for four hours. Cells were collected and stained with Ghost Dye Violet 450 (Tonbo Biosciences) followed by surface staining. Cells were then washed, fixed at room temperature for 15 minutes (PBS, 1% neutral buffered formalin), permeabilized and stained with antibodies against cytokines on ice (PBS, 2% FBS, 0.2% saponin). All samples were acquired on LSRFortessa or FACSVerse (BD Bioscience) and analyzed using FlowJo software V10.

Antibodies to the following were purchased from Biolegend: CD8 (53-6.7), IFNγ (XMG1.2), Epcam (G8.8); from eBioscience: IL17A (eBio17B7), IL13 (eBio13A), MHCII (I-A) (NIMR-4), Qa2 (69H1-9-9); from Tonbo Biosciences: CD4 (GK1.5), MHCII (I-A/I-E) (M5/114.15.2); The CD16/CD32 (2.4G2) antibody was obtained from UCSF’s Monoclonal Antibody Core.

### Thymic epithelial cell isolation

Thymic epithelial cells were isolated as described^7^. Briefly, thymi from 4-week-old mice were minced with razor blades into 1-2 mm^3^ pieces. The tissue fragments were collected and enzymatically digested three times (DMEM, 2% FBS, 100 μg/ml DNase I and 100 μg/ml Liberase™). The single-cell suspensions from each digestion were pooled into 20 ml of MACS buffer (PBS, 0.5% BSA, 2 mM EDTA) on ice. Total cells were washed and centrifugated using a three-layer Percoll gradient with specific gravities of 1.115, 1.065 and 1.0. Thymic stromal cells were enriched and collected from the Percoll-light fraction, between the 1.065 and 1.0 layers. Thymic stromal cells were washed, stained and sorted to isolate MHC-II^high^CD80^high^ mTEC (mTEC^high^) by FACS.

### RNA isolation and quantitative real-time PCR analysis

RNA was isolated with the EZNA Total RNA kit (Omega Bio-tek) was reverse transcribed to cDNA with SuperScript III Reverse Transcriptase and oligo d(T)16 primers (Invitrogen). Quantitative real-time PCR was performed on Bio-Rad CFX thermal cyclers with TaqMan Gene Expression assays from Life Technologies *(GAPDH,* Hs02786624_g1; *IFNB1,* Hs01077958_s1; *IFI6,* Hs00242571_m1; *IFI27,* Hs01086373_g1; *IFI44L,* Hs00915292_m1; *ISG15,* Hs01921425_s1; *IFIT1,* Hs03027069_s1; *IFITM1,* Hs00705137_s1; *RSAD2,* Hs00369813_m1; *Gapdh,* Mm99999915_g1; *Ifnb1,* Mm00439552_s1; *Ifit1,* Mm00515153_m1; *Ifi44l,* Mm00518988_m1; *Isg15,* Mm01705338_s1; *Rsad2,* Mm00491265_m1; *S1pr1:* Mm00514644).

### RNA-Seq and data analysis

RNA-sequencing libraries by first using the Nugen Ovation method (Tecan 7102-A01) to create cDNA from the isolated RNA, the sequencing library was then created from cDNA using the Illumina Nextera XT method (Illumina FC-131-1096). All libraries were combined and sequenced on Illumina HiSeq4000 lanes, yielding approximately 300 million single end, 50bp reads. Sequencing reads were then aligned to the mouse reference genome and the ensemble annotation build (GRCm38.78) using STAR^8^ (v2.4.2a). Read counts per gene were used as input to DESeq2^9^ (v1.26.0) to test for differential gene expression between conditions using a Wald test. Genes passing a multiple testing correct p-value of 0.1 (FDR method) were considered significant. Pathway analysis was carried out using DAVID^10^ and the R/Bioconductor package RDAVIDWebService^11^ (v1.24.0).

### Statistical analysis

All statistical analysis was performed using Prism 7 (GraphPad Software) or R 4.0.0 (R Foundation for Statistical Computing). Where indicated, two-tailed Mann-Whitney *U-*test and unpaired, parametric, two-tailed student’s *t*-test were used to evaluate the statistical significance between two groups, and *p* <0.05 was considered statistically significant.

## Acknowledgements

Suzy A. A. Comhair and Serpil C. Erzurum from Cleveland Clinic Foundation for help providing lung explants. Paul Wolters, Tien Peng, Chaoqun Wang and N. Soledad Reyes de Mochel at UCSF for help with providing lung explants. David J. Erle for discussions and assistance with RNA-sequencing. Mark S. Anderson for discussions and review.

## Author contributions

ZD and ZC designed and performed experiments, analyzed data. KM and TT provided technical advice. CSL performed experiments and analyzed data. FOH performed experiments. BJB assisted with small molecule experimental design. WLE analyzed RNA-sequencing data. TM provided lung explant tissue. AKS directed the study and wrote the manuscript.

## Data and materials availability

All data that support the findings of this study are available from the corresponding author upon reasonable request. RNA-sequencing data will be uploaded into a public respository and accession codes and unique identifiers will be provided.

## Extended Data

**Extended Data Fig. 1.**
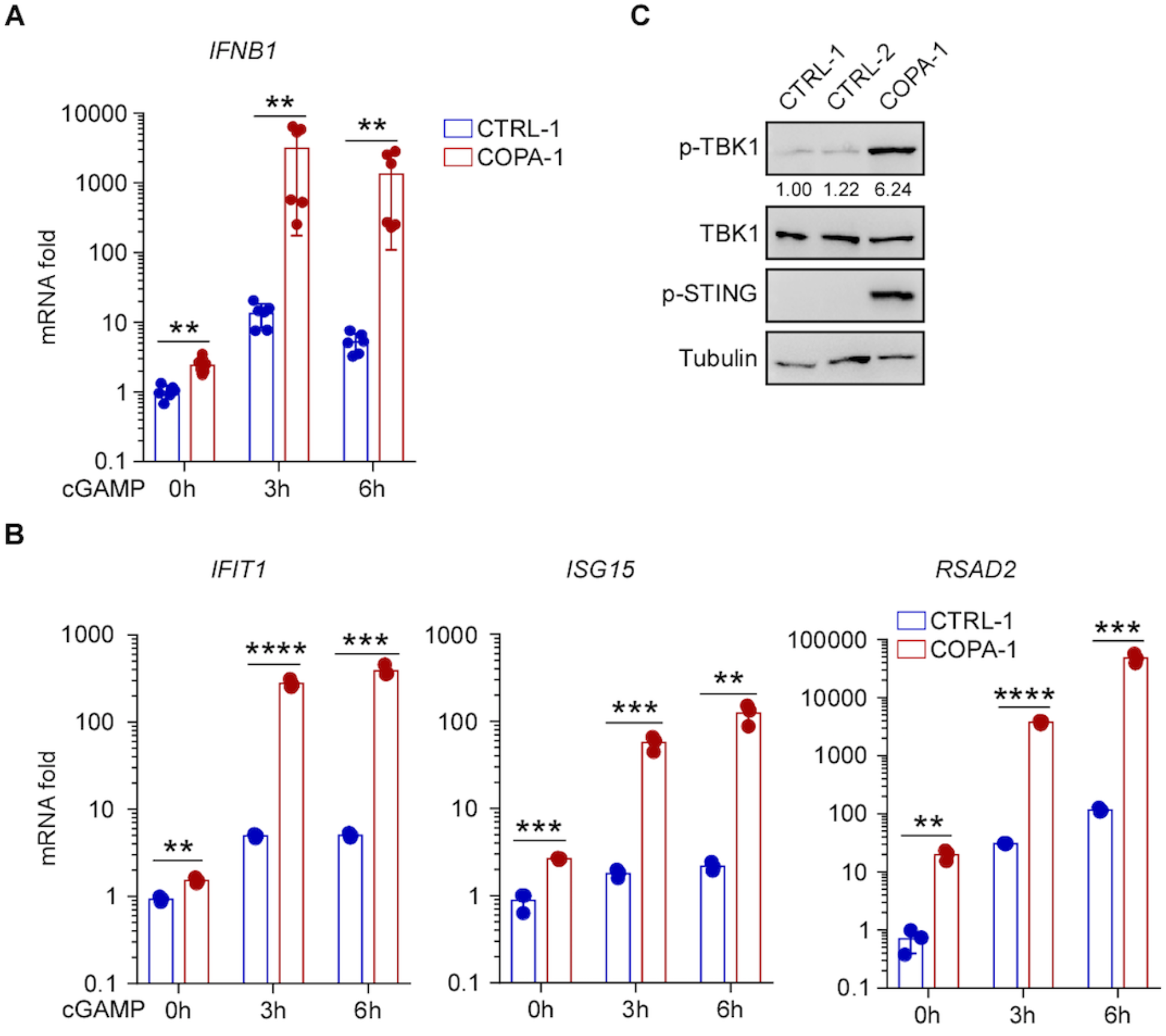
COPA syndrome patient fibroblasts demonstrate STING activation. (**A**) Real-time PCR was performed for *IFNB1* expression in primary lung fibroblasts from a healthy control and COPA syndrome subject treated with cGAMP (2μg/ml) for indicated time. Data pooled from 2 independent experiments are means ± SD, two-tailed Mann-Whitney *U* test. (**B**) Real-time PCR for ISG expression in primary lung fibroblasts from a healthy control and COPA syndrome subject treated with cGAMP (2μg/ml) for indicated time. Data are means ± SD, unpaired two-tailed Student’s *t* test. (**C**) Immunoblots of primary lung fibroblasts for indicated antibodies. **p<0.01, ***p<0.001, ****p<0.0001.

**Extended Data Fig. 2.**
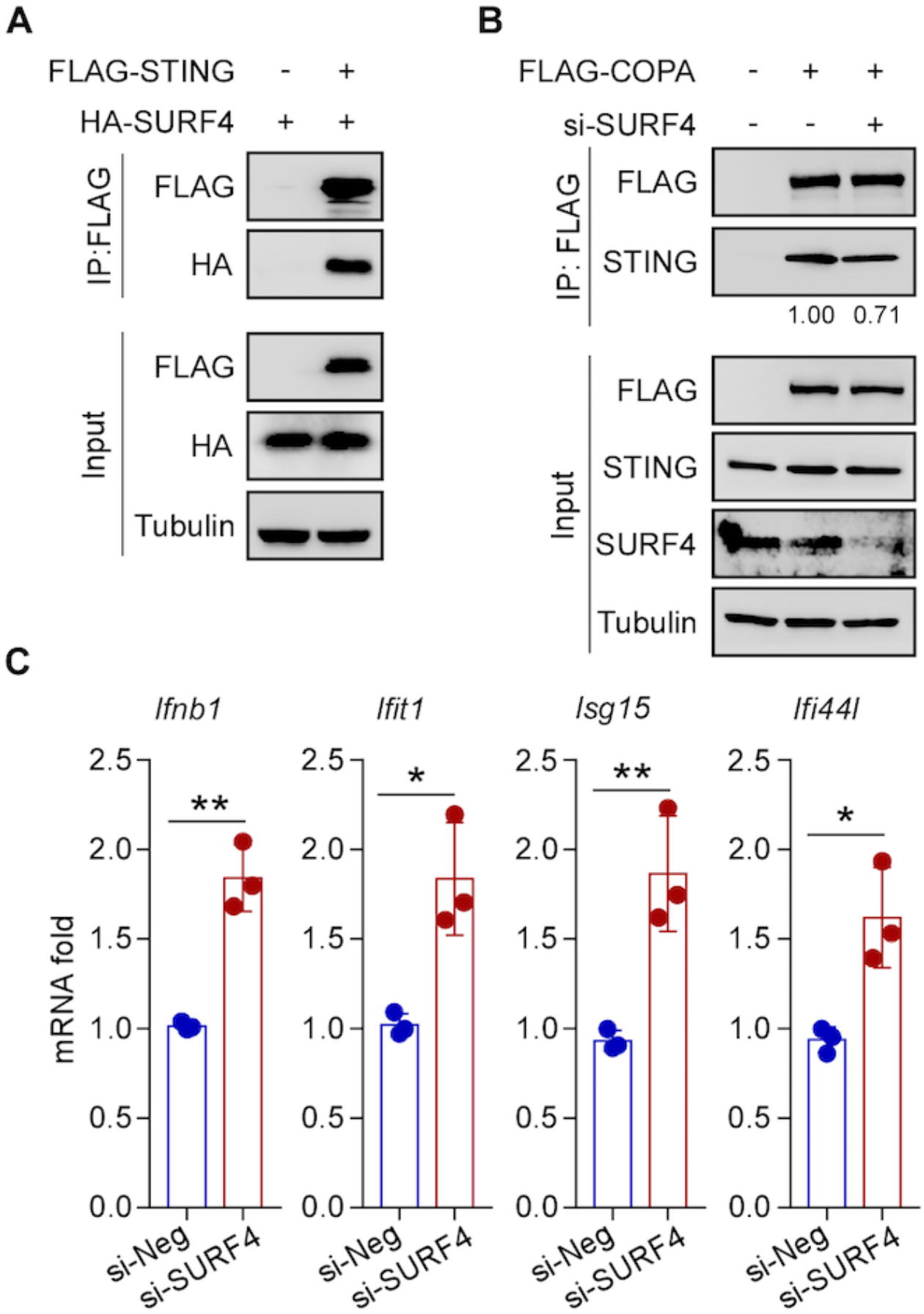
SURF4 functions as a cargo receptor that mediates retrieval of STING by COPA. (**A**) FLAG immunoprecipitates (IP) from lysates of HEK293T cells overexpressing FLAG-tagged STING and HA-tagged SURF4 were immunoblotted for indicated antibodies. (**B**) HEK293 cells were transfected with FLAG-tagged COPA and si-SURF4 for 48 hours. FLAG immunoprecipitates (IP) from lysates of HEK293 cells were immunoblotted for indicated antibodies. (**C**) Real-time PCR for ISG expression in immortalized MEF cells with EGFP-STING after si-SURF4 for 48 hours. Data represent means ± SD. *p<0.05, **p<0.01 (unpaired two-tailed Student’s *t* tests). siNeg = non-targeting siRNA control.

**Extended Data Fig. 3.**
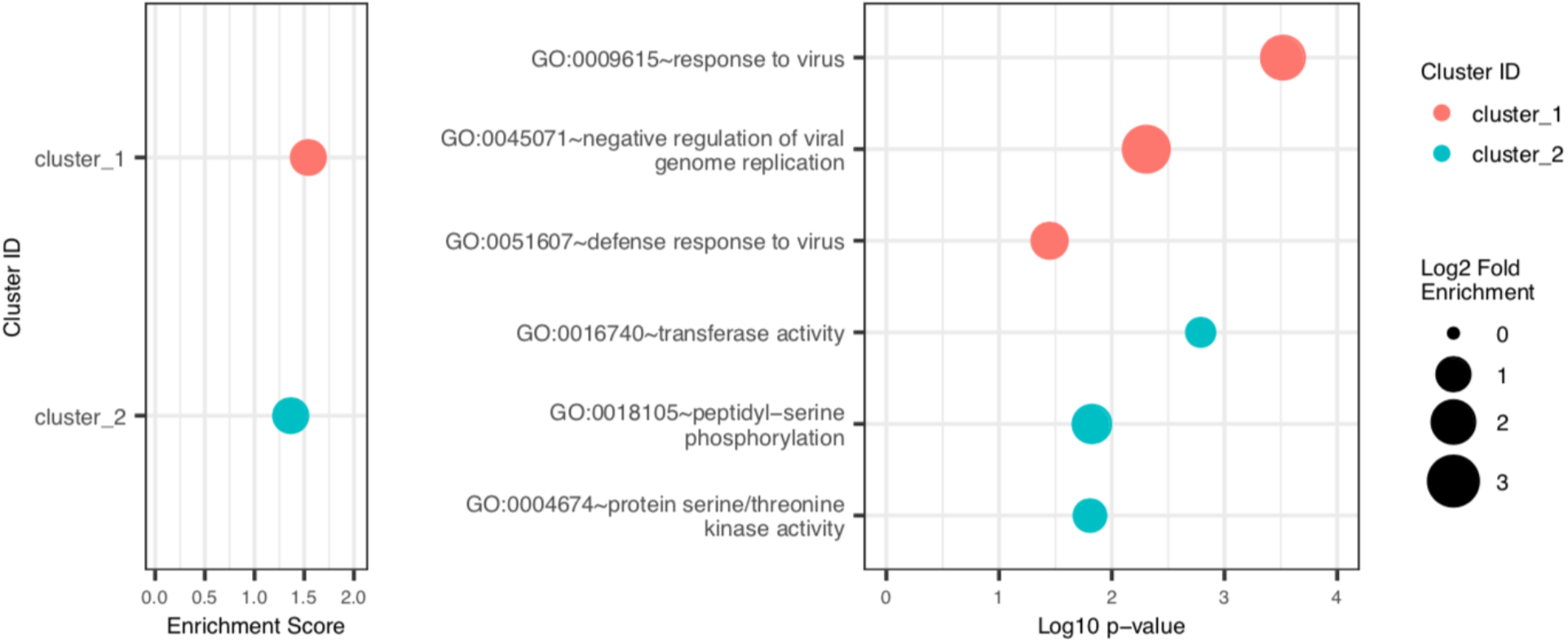
DAVID Gene Ontology analysis of bulk RNA sequencing in thymic epithelial cells. DAVID Gene Ontology analysis comparing differentially expressed genes in MHC-II^high^CD80^high^ mTECs from WT and *COPA^E24IK/+^* mice. Top GO clusters with the key word are shown. (n=3 of each genotype).

**Extended Data Fig. 4.**
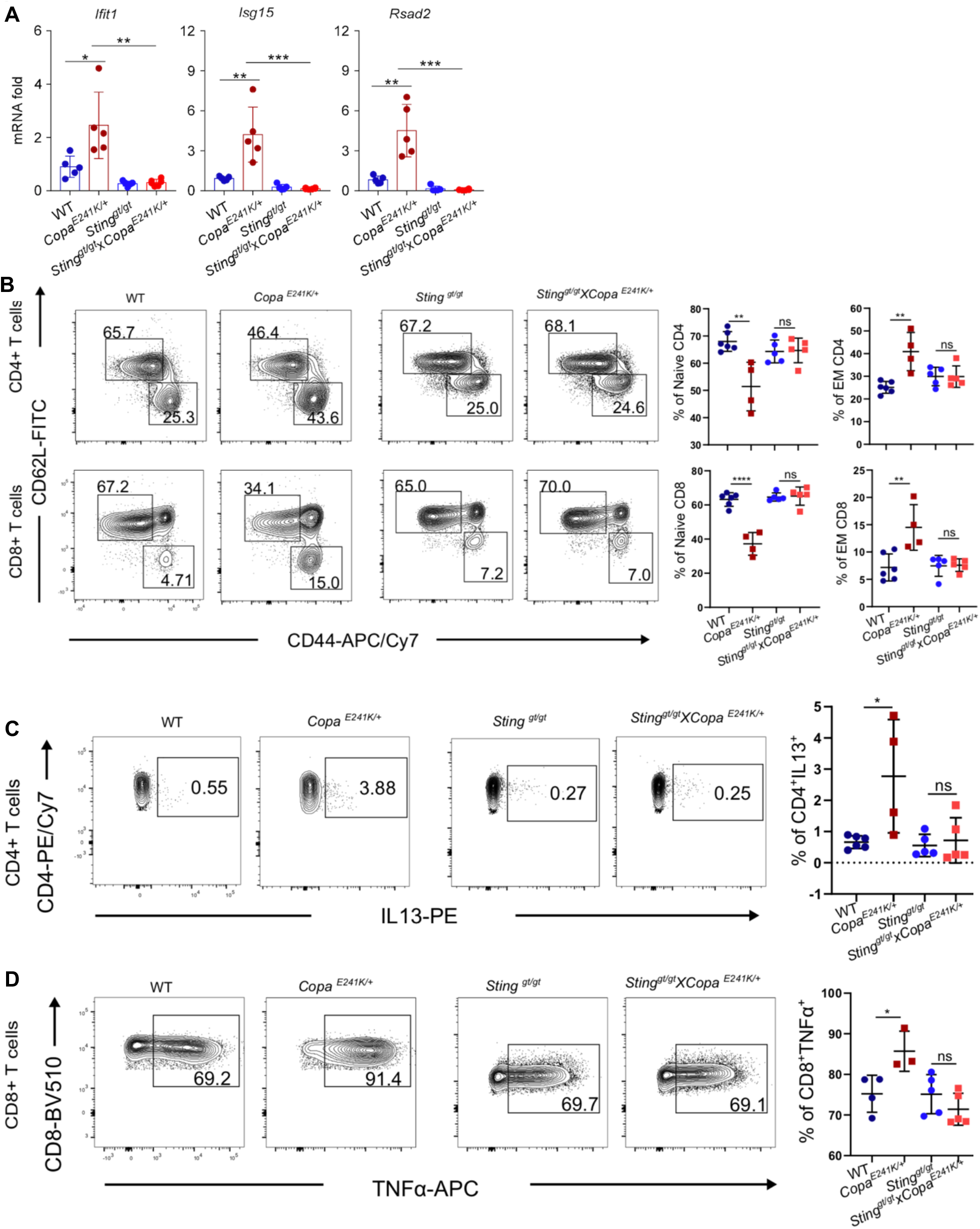
Activated STING in *Copa^E241K/+^* mice results in type I interferon driven inflammation. (**A**) Real-time PCR performed for ISG expression in splenocytes from WT (n=5), *Copa^E241K/+^* (n=5), *Sting^gt/gt^* (n=5) and *Copa^E241K/+^xSting^gt/gt^* (n=6) mice. (**B**) left: Representative flow plots showing expression of CD62L versus CD44 on T cells (top: CD4^+^ T cells; bottom: CD8^+^ T cells). right: Percentages of naive and effector memory T cells (left: CD4^+^ T cells; right: CD8^+^ T cells). (WT: n=6; *Copa^E241K/+^:* n=4; *Sting^gt/gt^*: n=5; *Copa^E241K/+^xSting^gt/gt^*: n = 5). (**C**) Left: intracellular levels of IL13 in splenic CD4^+^ T cells after PMA/Ionomycin stimulation. right: percentages of IL13 producing CD4^+^ T cells. (WT: n=6; *Copa^E241K/+^*: n=4; *Sting^gt/gt^*: n=5; *Copa^E241K/+^xSting^gt/gt^:* n=5, > 3 independent experiments). (**D**) Left: intracellular TNFα production in splenic CD8^+^ T cells after PMA/Ionomycin stimulation. right: percentages of TNFα producing CD8^+^ T cells. (WT: n=4; *Copa^E241K/+^*: n=3; *Sting^gt/gt^:* n=5; *Copa^E241K/+^xSting^gt/gt^*: n=5, >3 independent experiments). Data in (A-D) represent means ± SD. *p<0.05, **p<0.01, ****p<0.0001 (unpaired two-tailed Student’s t tests).

**Extended Table 1:** STING interacting partners containing a C-terminal dilysine motif. Seven STING interacting partners with C-terminal dilysine motifs were identified in published studies describing STING affinity purification-mass spectrometry (AP-MS) data^13–16,27^. Shown for each protein are its C-terminus sequence with dilysine motif highlighted, subcellular location and function.

| Protein | C terminus | Subcellular location | Function |
| --- | --- | --- | --- |
| CCDC47 | KQIKV<u>K</u>AM | ER | Calcium regulation protein |
| DDOST | MKE<u>K</u>E<u>K</u>SD | ER | Glycosyltransferase enzyme |
| HM13 | KGLE<u>K</u>KEK | ER, Plasma membrane | ER-resident intramembrane protease |
| MATR3 | ERRQ<u>K</u>KET | Nucleus | RNA binding protein |
| SAC1 | LVQ<u>K</u>E<u>K</u>ID | ER, Golgi | Phosphoinositide phosphatase |
| STT3B | KKIS<u>K</u>KTV | ER | Catalytic subunit of oligosaccharyl transferase |
| **SURF4** | **MDEK<u>K</u>KEW** | **ER, ERGIC, Golgi** | **Cargo receptor** |

**Extended Table 2.** Loss of STING rescues embryonic lethality of *Copa^E241K/E241K^* mice.

|  | *Sting*<sup>wt/wt</sup> | *Sting*<sup>gt/gt</sup> |
| --- | --- | --- |
|  | Neonates from 5 litters | Neonates from 5 litters |
| *Copa*<sup>+/+</sup> | 7 (24%) | 11(22%) |
| *Copa*<sup>E241K/+</sup> | 22 (76%) | 22(44%) |
| *Copa*<sup>E241K/E241K</sup> | 0 (0%) | 17 (34%) |

